# Pan-Cancer Analysis Reveals Technical Artifacts in TCGA Germline Variant Calls

**DOI:** 10.1101/092163

**Authors:** Alexandra R. Buckley, Kristopher A. Standish, Kunal Bhutani, Trey Ideker, Hannah Carter, Olivier Harismendy, Nicholas J. Schork

## Abstract

The degree to which germline variation drives cancer development and shapes tumor phenotypes remains largely unexplored, possibly due to a lack of large scale publicly available germline data for a cancer cohort. Here we called germline variants on 9,618 cases from The Cancer Genome Atlas (TCGA) database representing 31 cancer types. We identified batch effects affecting loss of function (LOF) variant calls that can be traced back to differences in the way the sequence data were generated both within and across cancer types. Overall, LOF indel calls were more sensitive to technical artifacts than LOF Single Nucleotide Variant (SNV) calls. In particular, whole genome amplification of DNA prior to sequencing led to an artificially increased burden of LOF indel calls, which confounded association analyses relating germline variants to tumor type despite stringent indel filtering strategies. Due to the inherent noise we chose to remove all 614 amplified DNA samples, including all acute myeloid leukemia and virtually all ovarian cancer samples, from the final dataset. This study demonstrates how insufficient quality control can lead to false positive germline-tumor type associations and draws attention to the need to be sensitive to problems associated with a lack of uniformity in data generation in TCGA data.

**Author Summary:** Cancer research to date has largely focused on genetic aberrations specific to tumor tissue. In contrast, the degree to which germline, or inherited, variation contributes to tumorigenesis remains unclear, possibly due to a lack of accessible germline variant data. In this study we identify germline variants in 9,618 samples using raw germline exome data from The Cancer Genome Atlas (TCGA). There are substantial differences in the way exome sequence data was generated both across and within cancer types in TCGA. We observe that differences in sequence data generation introduced batch effects, or variation that is due to technical factors not true biological variation, in our variant data. Most notably, we observe that amplification of DNA prior to sequencing resulted in an excess of predicted damaging indel variants. We show how these batch effects can confound germline association analyses if not properly addressed. Our study highlights the difficulties of working with large public genomic datasets like TCGA where samples are collected over time and across data centers, and particularly cautions the use of amplified DNA samples for genetic association analyses.

## Introduction

Cancer research to date has largely focused on genetic aberrations that occur specifically in tumor tissue. This is not without reason as tumor formation is driven to a great degree by somatically-acquired changes[1]. However, the degree to which germline, or inherited, DNA contributes to tumorigenesis is unknown. While it has been clearly demonstrated that germline variation increases cancer risk in overt and rare familial cancer predisposition syndromes, the contribution of germline to more common and sporadic cancer risk is unclear and highly debated[1,2]. It is likely that inherited germline variation in fundamental molecular processes, such as DNA repair, can create a more permissive environment for tumorigenesis and shape tumor growth in some individuals.[3
–5]. It is also likely that variation in the host germline genome can act synergistically with acquired somatic mutations to shape the way in which tumors grow and ultimately manifest.

There is a growing interest in better understanding the contribution of germline variation to cancer risk and tumor phenotypes[6,7]. The most extensive pan-cancer germline study to date identified associations between deleterious germline variation in known cancer predisposing genes and both age of onset and somatic mutation burden[6]. This study demonstrates that inherited variants can increase risk of developing cancer, as well as influence tumor growth and overall phenotypic features. Similar results were found in a study of bialleleic mismatch repair deficiency (bMMRD). It is known that bMMRD predisposes to childhood cancer, but it was further demonstrated that acquisition of somatic mutations in polymerase genes *(POLE, POLD1)* led to a hypermutated phenotype in childhood brain tumors[8]. This demonstrates a synergistic interaction between germline variation and somatic mutation. A comprehensive study of breast cancer whole genomes identified a somatic copy number profile signature associated with *BRCA1* inactivation[9]. Interestingly, this profile was associated with either inactivation of *BRCA1* in the tumor via mutation or promoter hypermethylation, or via inherited germline variants. This shows that somatic mutation and germline variation can both influence tumor phenotype.

We chose to use the whole exome sequence (WXS) data from TCGA to investigate the role of germline variation in shaping tumor phenotypes. TCGA is an attractive dataset for this purpose as there are paired tumor normal data for many cancer types. We took a pan-cancer approach for two reasons: 1. increased sample size and therefore increased power to detect associations of small effect size; and 2. cancers of disparate origin may share common features which would be overlooked in a cancer type-specific analysis[10]. For example, germline mutations in BRCA1/2 are most commonly studied in breast and ovarian cancer, but have also been shown to increase risk for stomach and prostrate cancer[11]. Further, germline BRCA2 mutations have been associated with a distinct somatic mutational phenotype and an overall increased somatic mutation burden in both prostrate and breast cancer[6,9,12]. To our knowledge, a comprehensive germline analysis of all cancer types available in TCGA has not been performed. Thus other cross-cancer germline associations likely remain to be discovered.

In an ideal dataset, a single protocol should be used for processing all samples. Unfortunately, this is unrealistic in large public datasets like TCGA in which samples are collected over time and across many data centers. Since its inception in 2005, TCGA has collected data on 11,000 patients from 20 collaborating institutions and generated sequence data from 3 sequencing centers[13]. Differences in sample collection and processing across centers could lead to batch effects, or variation in the data due to a technical factor that masks relevant biological variation[14]. Problems with batch effects can be magnified when analyzing samples across TCGA, since the number of methods used to collect samples increases with the number of cancer types. The Pan-Cancer Analysis Project has recognized this and aims to generate a high quality dataset of 12 TCGA cancer types, taking care to identify and minimize technical artifacts[10].

While extensive curated somatic data is available from TCGA, germline information is currently only available in raw form, under controlled access. Therefore, we first had to develop and execute a variant calling pipeline on the raw normal tissue sequence data to generate TCGA germline variant calls. As a main goal of our variant calling analysis is to create a cohesive, pan-cancer dataset, we chose to use the Genome Analysis Toolkit (GATK) group-calling approach[15,16]. Group-calling is a strategy for variant calling in which read data is shared across samples, in contrast to single sample calling where genotype decisions are made based on reads from a single sample only. There are three major advantages of this approach: the ability to distinguish sites that are homozygous reference vs. those that have insufficient data to make a call, increased sensitivity to detect variant sites that are poorly covered in any individual sample but well covered when the cohort is considered as a whole, and the ability to use GATK's statistical modeling approach to variation filtration, known as ‘variant quality score recalibration’ (VQSR).

Here we describe our experience calling germline variants from a large cohort of TCGA normal tissue WXS samples spanning 31 cancer types. Specifically, we were interested in cataloguing sources of heterogeneity in sample preparation, identifying batch effects in our variant calls, and determining methods to reduce or control for technical noise. Our findings highlight the importance of quality control at all stages of the variant calling process and suggest that pan-cancer analysis with TCGA data be approached with caution.

## Results

### Technical heterogeneity in TCGA WXS Data Generation

We obtained TCGA WXS data from CGhub in the form of reads aligned to the human reference genome (BAM files)[17]. From the BAM files and available metadata we identified seven technical sources of variation in the way the sequence data were generated: tissue source of normal DNA, exome capture kit, whole genome amplification of DNA prior to sequencing (WGA), sequencing center, sequencing technology, BWA version, and capture efficiency (C20X) (S1 Fig). We found substantial variation existed within and between cancer types with respect to these technical factors (Fig 1). Some of these technical factors were found to be highly associated with cancer type, such as use of Illumina Genome Analyzer II and ovarian cancer (OV), while others exhibited no clear relationship with cancer type, such as use of solid normal tissue as opposed to blood as a source of normal DNA. Relationships existed between pairs of technical factors as well, such as the Broad Institute's exclusive use of a custom Agilent exome capture kit. All possible combinations of the first six technical factors produce 1152 unique workflows, of which only 44 were used to generate the TCGA data. This further demonstrates that relationships exist between technical factors. Of the 31 cancer types examined, only uveal melanoma (UVM) and testicular germ cell tumors (TCGT) had a uniform workflow for all samples (S2 Fig). These observations highlight the substantial heterogeneity in data generation across TCGA and importantly even within cancer types.

**Fig 1.**
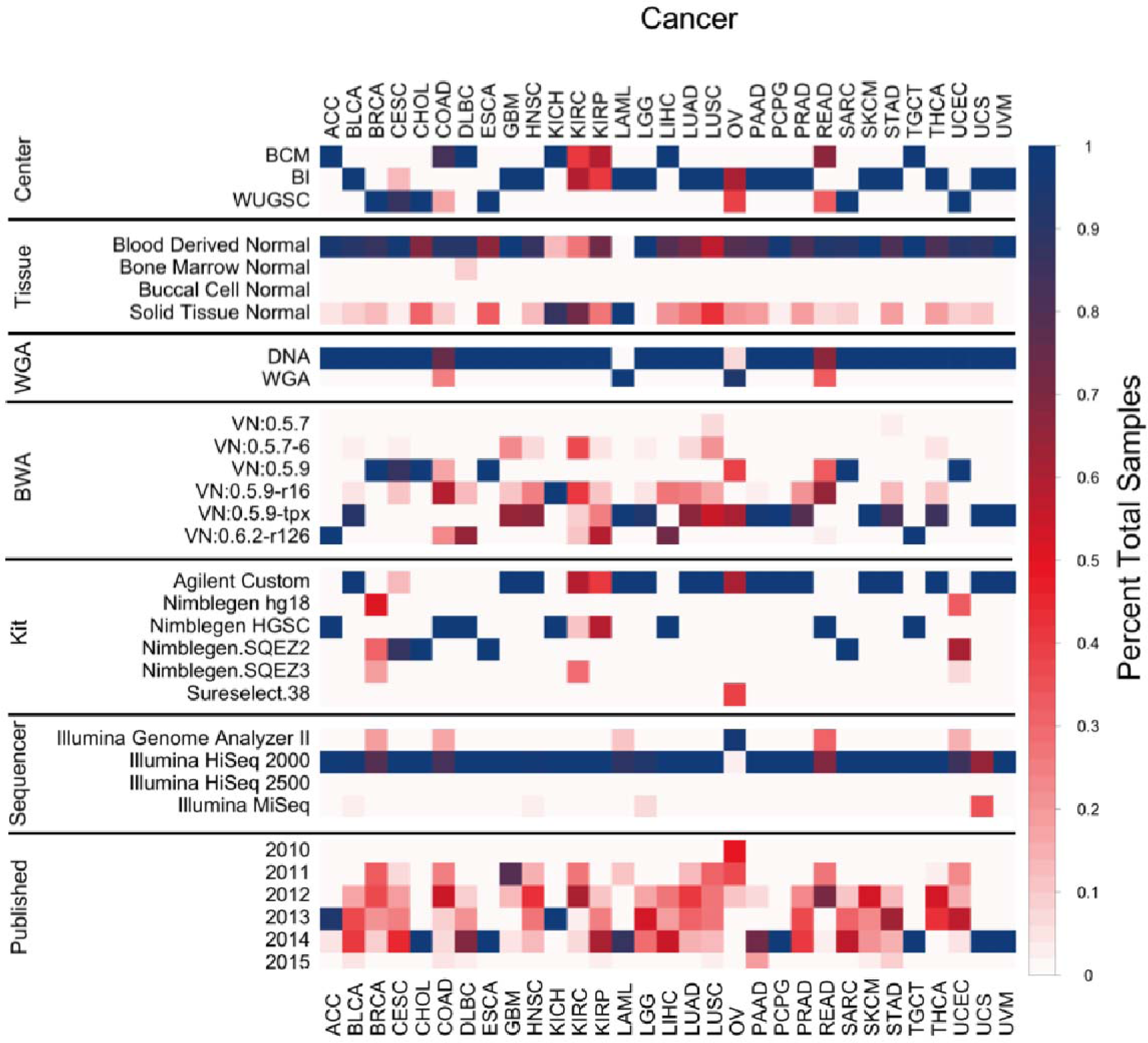
Overview of technical covariates for pan-cancer samples. For each covariate and cancer type, color represents the fraction of total samples. Fraction of total samples sums to 1 for each covariate and cancer type. Red indicates higher heterogeneity. Year first published included for context.

The technical factors can ultimately be divided into two categories: those that can be modified during processing of the sequence data (BWA version, target regions of a capture kit), and those that cannot be modified computationally (source of normal DNA, WGA, center, technology, capture efficiency). Six exome capture kits ranging in size from 33-64 MB were used to capture normal DNA for sequencing (S2 Table). As the goal of our variant calling pipeline was obtain a uniform set of variants across samples, we chose to limit analysis to the common capture regions. The area lost by confining attention to the intersection region of the capture kits is largely in exon flanking regions. The intersection covers 97.7% of Gencode exons, thus for the purposes of studying protein-coding variation using the intersection of the kits leads to minimal loss of data (S2 Table)[18]. It has been shown that differences in capture efficiency and sample preparation protocols between exome kits can affect variant calls, even in regions common between kits[19]. Therefore, despite using the common capture region, the use of multiple capture kits may still introduce artifacts.

To assess the effect of heterogeneous BWA alignments on variant calls, we called variants on 345 of the TCGA normal WXS samples either using the provided BAM (OldAlign) or stripping and realigning reads to GRCh37 using BWA MEM v.0.7.12 (NewAlign) (S3 Fig). The overall discordance rates between the two sets of variants is in the expected range for different alignment protocols (S4 Fig)[20]. Indel calls were noticeably more discordant, consistent with the specific challenges and notorious variability of indel calling[21]. Greater discordance between variant calling pipelines has been observed in low complexity regions of the genome, and in accordance with this we observe much lower discordance rates after removing repetitive regions from analysis (S4 Fig)[22]. Interestingly, the discordance rate was correlated with BWA version used to generate the BAM file in CGhub, with older versions displaying more discordance. This effect can largely be reduced by filtering the variants (S5 Fig). In the absence of a truth set of variants we cannot determine whether realigning BAM files produces more accurate calls. Given the computational cost of realignment, and that discordance can be mitigated by filtering variants and masking repetitive regions of the genome, we proceeded with variant calling using the provided BAM files.

Functional annotation of the final VCF predicted 1,093,501 variants (625,365 missense; 371,754 silent; 24,455 nonsense; 2,968 splice site; 553 stoploss; 46,280 frameshift indels and 22,126 in-frame indels) in 9,618 samples. For initial quality control we performed principal component analysis (PCA) to identify the most significant sources of variation in the variant calls. PCA on common variants showed that the first two principal components stratified samples by self-reported race and ethnicity, indicating that the largest source of variation is ethnic background and not technical factors (S6 Fig). To assess the quality of the calls, we measured the fraction of variants also present in the ExAC database[23]. We expect a high degree of overlap between our calls and ExAC, as the ExAC samples include 7,601 normal DNA samples from TCGA. Overall 88.56% of the variant calls were present in ExAC, with SNVs showing higher overlap than indels (89.91% vs. 53.94%). Taken together, we concluded the variant calls were free of overt technical artifacts and proceeded to the next stage of analysis.

### Impact of technical heterogeneity on loss of function variants

We are ultimately interested in testing the hypothesis that inherited impaired functionality of cancer-relevant pathways shape tumor phenotypes, as has been previously demonstrated for bMMRD and BRCA1 germline mutations[6,8,9]. To identify germline variation likely to disrupt function of genes, we used VEP and LOFTEE to predict LOF variants in this cohort[24]. We observed a median 150 LOF per sample across our entire cohort, consistent with the ExAC findings (Fig 2). [23]. However, two cancer types, acute myeloid leukemia (LAML) and OV deviate significantly from this expected value, with individuals with these cancers having up to 500 LOF germline variants. This suggests an artifact was manifesting in rare LOF variants that was not identified by PCA on common variants. Notably this effect is specific to LOF indels, in contrast to LOF SNVs that are distributed more uniformly across cancer type (S7 Fig).

**Fig 2.**
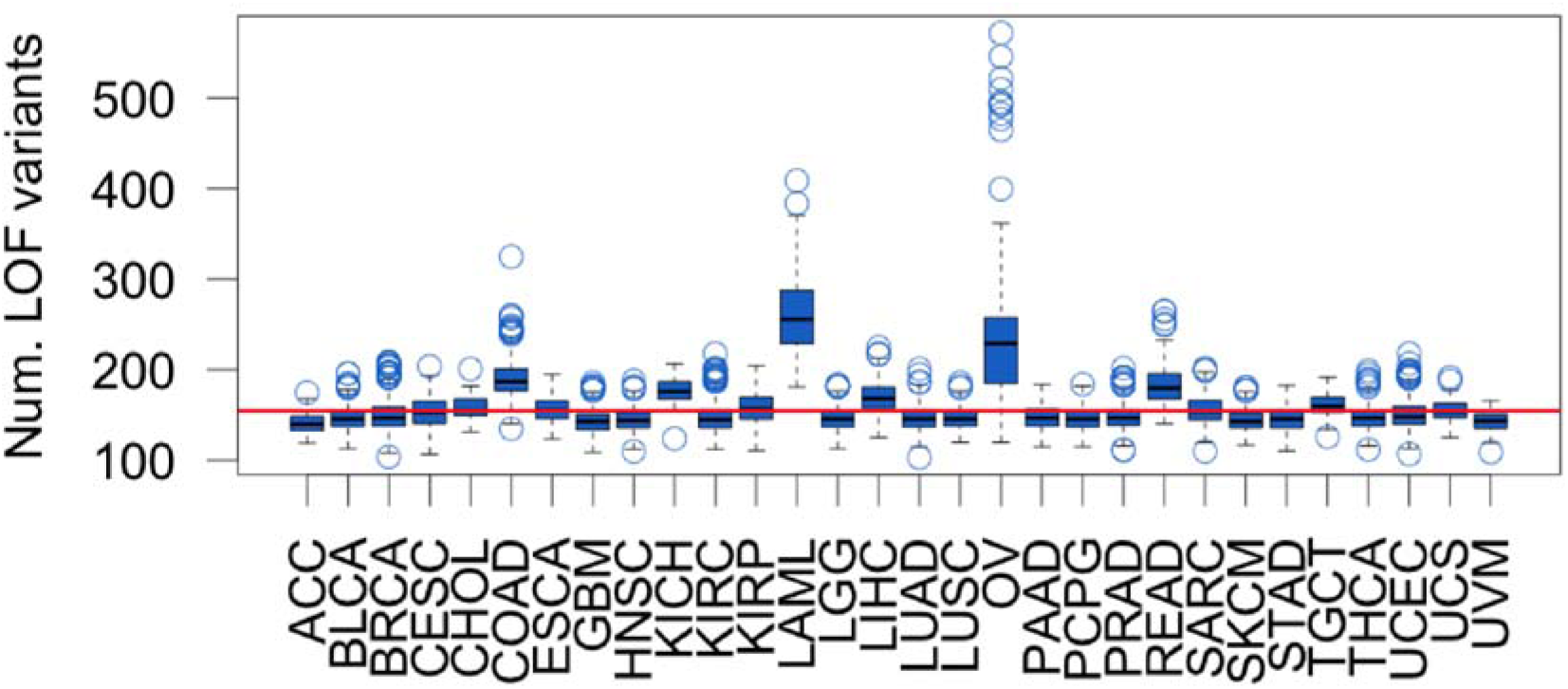
Individual LOF variant burden across cancer types. LOF variant burden includes both SNV and indels. Red line indicates expected LOF burden from ExAC (155).

We used Analysis of Variance (ANOVA) to assess the contribution of each technical factor to individual LOF variant burden. Initial analysis showed that source of normal control DNA and sequencing technology were not significantly associated with LOF variant burden, and that capture kit was highly collinear with sequencing center. Therefore, we limited subsequent analysis to sequencing center, BWA version, WGA, and C20X. It is known that LOF variant burden varies between ethnic groups, thus we include self-reported race as a covariate in this analysis as a reference point for expected variation[23]. All technical factors combined explain less than 1% of the variance in LOF SNV burden, indicating SNVs are largely unaffected by technical variation. In contrast, 59% of variation in LOF indel burden was explained by technical factors, with WGA alone explaining over 50% (Table 1).

**Table 1.** Variance in LOF SNV and indel burden explained by technical covariates.

| LOF SNV |  |  |  |  | LOF Indel |  |  |  |
| --- | --- | --- | --- | --- | --- | --- | --- | --- |
|  | Sum Sq | Df | F value | Pr(>F) | Sum Sq | Df | F value | Pr(>F) |
| **C20X** | 1785.49 | 1 | 52.95 | 3.72E-13 | 52930.43 | 1 | 153.90 | 4.95E-35 |
| **WGA** | 156.89 | 1 | 4.65 | 3.10E-02 | 3744887.28 | 1 | 10888.62 | 0.00E+00 |
| **Center** | 716.08 | 2 | 10.61 | 2.48E-05 | 383585.43 | 2 | 557.65 | 4.53E-228 |
| **BWA** | 79.59 | 5 | 0.47 | 7.97E-01 | 169507.9 | 5 | 98.57 | 2.76E-101 |
| **RACE** | 30698.90 | 5 | 182.10 | 1.33E-184 | 146904.86 | 5 | 85.42 | 7.59E-88 |
| **Residuals** | 281966.85 | 8363 |  |  | 2876257.21 | 8363 |  |  |

WGA samples have a higher LOF variant burden with a median 201 LOF variants per WGA sample. Four cancer types contain samples that underwent WGA: colon adenocarcinoma (COAD) (26% WGA), rectum adenocarcinoma (READ) (33% WGA), OV, (92% WGA) and LAML (100% WGA) (Fig 1). Analyzing cancer types containing both amplified and non-amplified DNA samples, we observed that WGA samples had a significantly higher LOF variant burden (Fig 3A), further suggesting that WGA rather than cancer type is the main source of bias. The cohort contains 13 individuals with both amplified and non-amplified DNA samples. We observed a 1.5 fold increase in LOF variant burden in amplified samples relative to non-amplified samples from the same individuals (p = 0.0002 by paired Wilcoxon Signed Rank test) (Fig 3B), suggesting that WGA prior to sequencing leads to an artificially inflated estimate of LOF variants.

**Fig 3.**
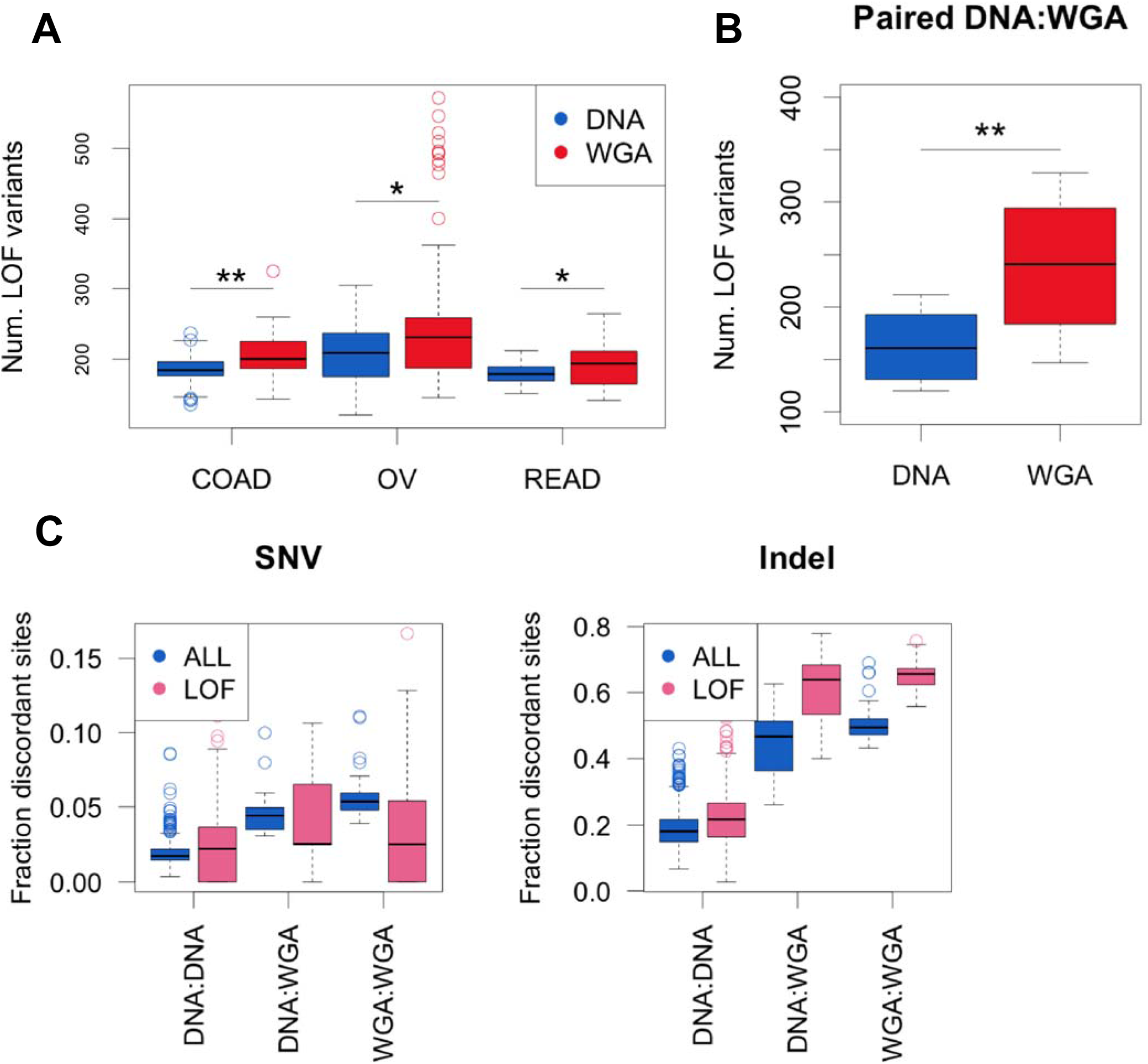
WGA increases LOF variant burden. (A) Individual LOF variant burden in cancers with WGA samples plotted by WGA status.. *= Wilcoxon rank sum test p < 0.05, ** = Wilcoxon rank sum test p < 0.001. (B) Individual LOF variant burden in N = 13 samples that have both DNA and WGA samples available. ** = Wilcoxon paired rank sum test p < 0.001. (C) Discordance between repeated samples of N = 493 individuals plotted by WGA status. DNA:DNA = all samples are DNA, WGA:DNA = at least one sample is WGA, WGA:WGA= all samples are WGA. Discordance was calculated separately for SNPs and indels using all variants or LOF predicted variants only.

Having demonstrated that WGA is associated with increased LOF variants, we sought to characterize WGA samples more deeply. We observe that WGA samples have an excess of LOF indels while LOF SNV burden appears unaffected, as expected from the ANOVA results (S8 Fig). Interestingly, WGA samples had fewer variants overall, due to poor coverage over the capture regions (S8 Fig). We next investigated the effect of WGA on variant call discordance between repeated samples. In addition to the 13 individuals with paired normal WXS samples with and without amplification (denoted WGA:DNA), 44 individuals have paired normal WXS samples where both samples have been amplified (denoted as WGA:WGA) and 435 are paired samples without amplification (denoted DNA:DNA). We calculated variant call discordance between all repeated samples for SNVs and indels separately and observed a stepwise increase in discordance with amplification of one or both samples. This effect was most apparent in LOF indels, with a median 65.6% LOF indel discordance between repeated WGA:WGA samples (Fig 3C). Notably, the high discordance between repeated WGA samples suggests that the errors introduced by the amplification process are not consistent between samples and supports the hypothesis that WGA introduces random errors in the sequence data that manifest as LOF indels during the variant calling process.

To determine whether our choice not to realign BAM files contributed to the observed WGA effect, we calculated LOF variant burden in our NewAlign and OldAlign cohort using the same protocol. Realignment of the sequence data with BWA MEM increased the number of LOF calls per individual but overall LOF burden was highly correlated (Pearson R^2^ = 0.95) (S9 Fig). WGA explained a significant amount of variance in LOF variant burden in both NewAlign and OldAlign samples (S9 Fig). Thus we can conclude that realignment does not remove WGA artifacts observed in our variant calling pipeline.

### Filtering artifactual LOF variant calls

We next sought to find an appropriate filter to remove artifactual LOF variant calls in WGA samples. As SNV calls were largely robust to technical artifacts, we focused on filtering indels specifically (S7 Fig). We used two strategies available from GATK: 1) Statistical model filtering using VQSR with increasing stringency cutoffs (99%, 95%, 90%), and 2) Heuristic filtering (Hardfilter) based on fixed thresholds (QD > 2, FS < 200, ReadPosRankSum > −20)[16]. The four filtering methods varied in stringency, resulting in a median individual LOF indel burden ranging from 53-98 across methods (Fig 4A and S10 Fig). All filters preserved the expected variation in LOF burden across ethnic backgrounds (S11 Fig). To assess technical artifacts, we performed an ANOVA analysis as described in Fig 2 for each filtering approach, including the initial filter (GATK VQSR 99) as a reference (Fig 4B). VQSR 90 and VQSR 95 reduced technical artifacts to a similar degree, whereas VQSR 99 and Hardfilters performed poorly (S11 Fig and S3 Table). Of note, when more stringent filtering approaches are applied the variance explained by self-reported race increased, indicating that when artificial variation introduced by technical factors is reduced true biological variation is revealed.

**Fig 4.**
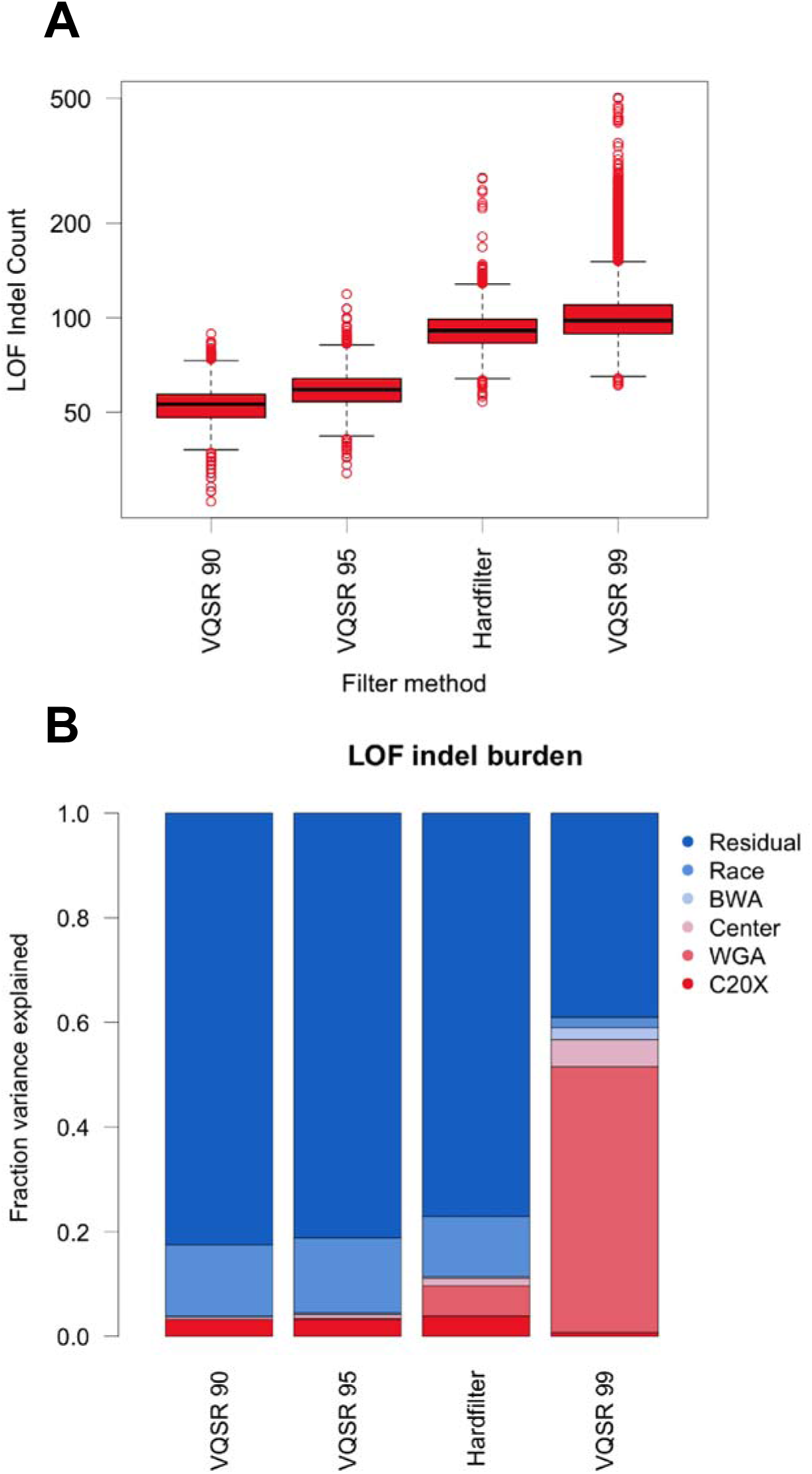
A comparison of indel filtering strategies. (A) Individual LOF indel burden for all indel filter methods in order of decreasing stringency. (B) Percent of variation in individual LOF indel burden explained by technical covariates for each filter method.

Variant filtering is a balance between removing likely false positive signal while retaining true positive signal. Using VQSR 99 we observe an individual LOF variant burden similar to that reported by ExAC, while all other methods produce lower LOF burden than expected (S12 Fig)[23]. Therefore, while more stringent filtering approaches can reduce technical artifacts, they do so at the cost of losing likely true positive indels. Without a way to manually validate a large number of rare indel variant calls, it is impossible to exactly measure false positives rates for our filter approaches. Instead, we used the repeated samples in our cohort to identify likely true positives (indels concordant between repeated samples) and likely false positives (indels discordant between repeated samples). We assessed filter quality using three measures: the fraction of discordant indels removed by the filter, the fraction of concordant indels removed by the filter, and the fraction of indels overlapping the ExAC database. The stringency of each filter was measured as the total number of LOF indel sites and the median individual indel LOF burden when each filter was applied (Table 2).

**Table 2.** Metrics of filter stringency and efficacy.

| Filter | LOF indel sites | Median LOF indel burden | Fraction discordant indels removed | Fraction concordant indels removed | Indel overlap with ExAC |
| --- | --- | --- | --- | --- | --- |
| VQSR 90 | 6212 | 53 | 0.8667 | 0.4514 | 0.7079 |
| VQSR 95 | 9177 | 59 | 0.8064 | 0.3760 | 0.6776 |
| Hardfilter | 24212 | 91 | 0.3600 | 0.0210 | 0.3527 |
| VQSR 99 | 26134 | 98 | 0.2763 | 0.1100 | 0.5394 |

By all quality measures GATK VQSR 90 performs the best, suggesting that high stringency is required to produce reliable indel calls.

Because it is possible that the observed WGA artifacts are specific to the GATK pipeline and that an alternate method of variant calling would reduce or eliminate this effect, we also obtained indel calls for all samples using Varscan (S13 Fig)[25]. While WGA artifacts are largely absent from Varscan indel calls, Varscan is highly sensitive to variation in BWA version. The Varscan pipeline does not include preprocessing or local indel realignment like the GATK pipeline, and likely would have performed better on realigned BAM files. It was not feasible to group call a cohort of this size using Varscan, and as we desired a group-called set of variants, we discontinued further analysis of Varscan indel calls.

### Consequences of technical artifacts on genetic associations

To determine how sensitive association results are to filtering method, we tested for association between germline LOF variant burden and cancer type using different filtering approaches. We took an 'one vs. rest' approach with our samples using all cancers except the cancer of interest as a control. Thus, we tested for enrichment of LOF germline variants in one cancer type as compared to other cancers, which is different than other studies that have used control cohorts[6]. Our rationale for using this approach was to minimize heterogeneity that would be introduced by including control samples collected in different studies. We chose to highlight the results only from OV for two reasons. First, it is established that *BRCA1/2* germline variants are enriched in OV so the OV – *BRCA1/2* association can be used as a positive control, and second virtually all OV samples have been amplified and are confounded with WGA artifacts[6,26,27].

QQ plots from logistic association tests for three indel filter methods are shown in Fig 5A. It was immediately apparent that our initial filtering approach (VQSR 99) produced an excess of significant associations even above a strict Bonferroni multiple hypothesis correction (Fig 5B). True associations are mixed with false associations due to WGA artifacts in LOF indel calls. Increasing the stringency of indel filtering reduced noise due to technical artifacts while retaining a putative true positive *BRCA1/2* association signal. Stringent filtering removes noise at the cost of reducing potential signal, as evidenced by the decreased number of genes that can be tested for association. This inflation in significant associations was only observed in WGA cancers, and persisted, albeit to a far lesser extent, even with the most stringent filter (Fig 5B). Supporting the idea that some of the associations in WGA cancer types are false is that fact that 25% of significantly associated genes were shared between LAML and OV. It is unlikely that two disparate cancer types would share a large percentage of germline associations for a biological reason. Further only two of the significant genes *(BRCA1/2)* in OV and none in LAML are genes where germline variation is known to be associated with cancer risk[28]. LOF indels calls are more prone to batch effects than LOF SNVs, therefore we repeated the association test limiting to LOF SNVs only. While this reduces the excess number of significant associations, the analysis was underpowered to detect the true positive *BRCA1/2* – OV association (S14 Fig). These results demonstrate that technical artifacts can lead to spurious associations and highlight the difficulty of correcting for artifacts in a pan-cancer analysis when technical factors are highly correlated with the phenotype being tested (Fig 1).

**Fig 5.**
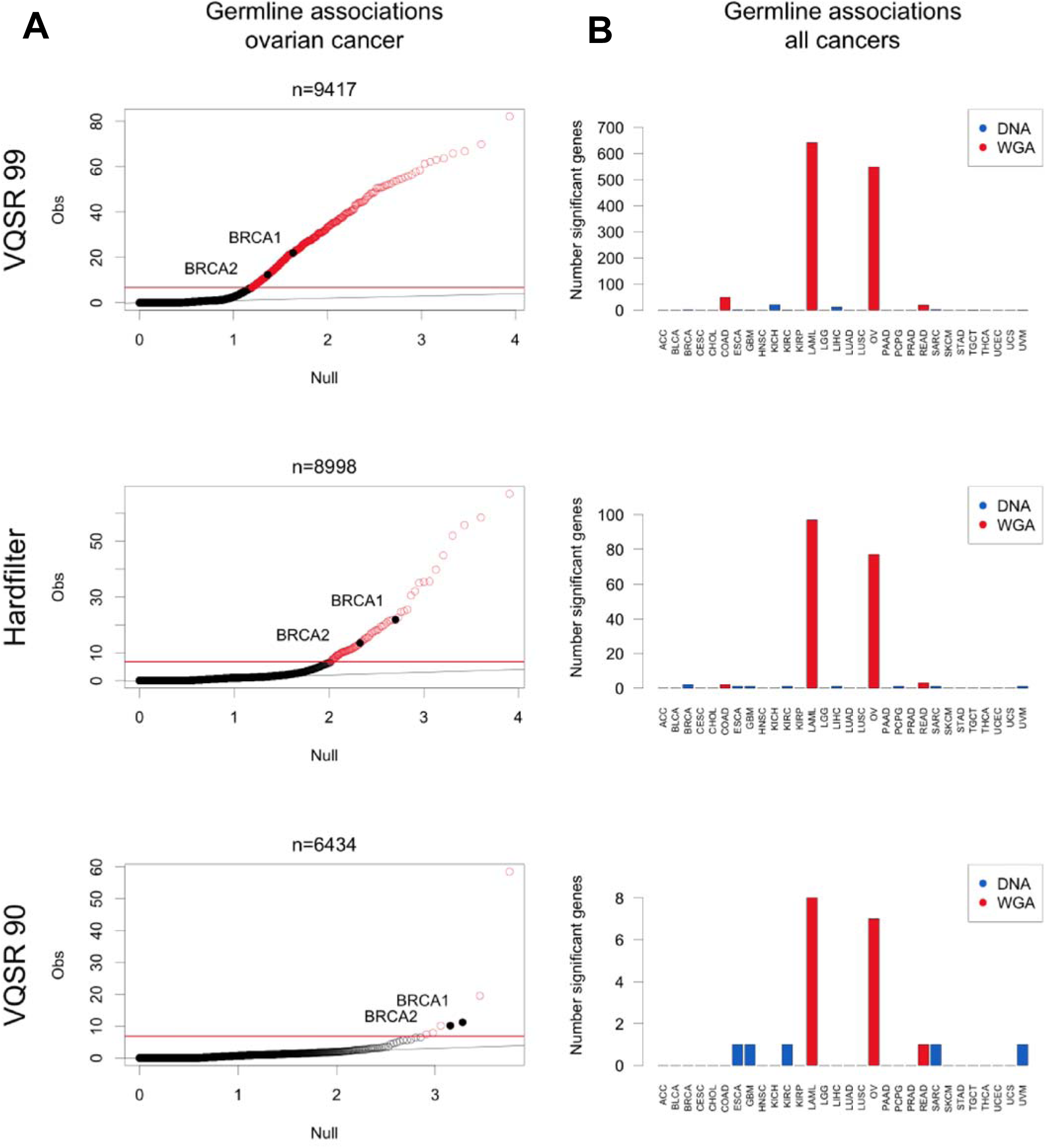
Association testing between germline LOF variant burden and cancer type. (A) QQ plots from logistic regression association testing between germline LOF burden and ovarian cancer for three indel filter methods. N = number of genes tested. Red indicates associations significant p < 1.61 × 10^−7^. *BRCA1/2* association highlighted. (B) Number of significant cancer type - gene associations in each cancer type for three indel filter methods. Color indicates cancer types with WGA samples.

We have demonstrated multiple problems that arise from including WGA samples in a pan-cancer cohort. Association tests revealed that despite stringent indel filtering, cancers with WGA samples have an excess of significant associations. The optimal indel filtering method to remove WGA artifactual signal removes approximately half of the expected LOF indels, and thus removes what is likely true signal from all samples. Finally, group calling shares information across all samples for both variant calling and filtering, so including samples with known artifacts may negatively affect all samples in the final VCF. Taken together, we decided to remove all WGA samples from future analysis, despite losing all LAML and samples and all but 30 OV samples. To properly remove WGA samples, we re-called variants beginning at the single sample gVCF stage. We compared the alleles found in the 9004 non-amplified DNA samples when called with the WGA samples (+WGA) and without (-WGA) and observed a surprising amount of discordance. This discordance was largely due to VQSR filtering, as the raw variant calls were highly concordant (S15 Fig). A similar magnitude of discordance was seen after removing a random subset of DNA samples and re-calling variants, indicating that this observation is not specific to WGA samples. From this we can conclude that group genotyping does not change dramatically with group size and membership, but VQSR is sensitive to group changes.

## Discussion

We identified sources of technical variation in LOF variant calls from TCGA germline WXS data. Overall SNV calls were more robust to technical factors than indel calls. We found the strongest association between amplification of DNA prior to sequencing and an excess of LOF indel calls. Other factors tested were found to be significantly associated with both LOF SNV and LOF Indel burden, but explain little of the total variance in LOF variant burden when appropriate filters are applied (Table 1 and Fig 4B). The factor explaining the most technical variation in total LOF variant calls after filtering is capture efficiency (C20X). It is likely that poor coverage over common capture regions, perhaps due to the different capture technologies used, decreased the ability to assign genotypes in some samples. Group-calling distinguishes sites with insufficient coverage to make a genotype call from those with adequate coverage for calling a homozygous reference genotype. Therefore, while C20X is a significant factor in the simple burden analyses performed here, a more sophisticated burden testing approach that can accommodate missing genotype values should mitigate this technical artifact.

Difficulty producing reliable variant calls in WGA exome samples has been previously reported[19,29]. Inaccurate read alignment has been identified as a main contributor to spurious calls in WGA samples. However, even with an alignment protocol optimized for WGA samples it is still estimated that 7% of variant calls in WGA samples are artifactual[19]. Artifacts introduced by WGA appear to be random and are therefore likely rare[19]. Previous work comparing amplified and nonamplified DNA obtained from the same biological sample report higher variant call discordance in indels compared to SNVs, similar to what we observe[29].These studies conclude that overall concordance between amplified and non-amplified samples is satisfactory; however, neither examined the impact of WGA on deleterious variants. Here we have demonstrated that random errors introduced by WGA tend to manifest as rare frameshift indels that are difficult to distinguish from true rare deleterious variation.

While group-calling is the approach recommended by GATK, with the exception of one paper from our lab exploring the impact of genetic background on group calling, to our knowledge there has not been a published systematic comparison of group calling vs. single sample calling on a gold standard dataset to quantify the advantages of group calling[30].Group-calling is not without problems. Greater accuracy for the group as a whole comes at the cost of loss of singleton variants from any given sample. Another complicating factor unique to group-called samples are multi-allelic sites, or sites where multiple alternate alleles are found in the population genotyped. Relatively few sites in our VCF were multi-allelic (3%, or 30,620 sites), but these sites contain 4,947 high-confidence LOF variants (11% of all LOF variants), indicating the importance of correct multi-allelic site parsing. Multi-allelic sites additionally pose a problem when filtering reliable from unreliable variants. With current tools for filtering VCFs, it is only possible to filter at the site level, meaning at multi-allelic sites all alleles will either be included or excluded by the filter. Further, in the version of GATK used for this analysis (v3.5), quality annotations for a site are calculated using all alternate reads without distinguishing between alleles. Therefore it is possible for low quality alternate alleles to pass filter at multi-allelic sites if high quality alternate alleles are present at the same site.

TCGA is often thought of as a single dataset, but due to differences in sample collection and processing across the participating sites, should be thought of as a collection of studies. While we focused on the germline WXS sequence data, it is likely that batch effects are present in other data types. This has been recognized by the Pan-Cancer TCGA effort, although it is less often acknowledged in papers published on one or few cancer types[10]. There is heterogeneity even within cancer types in terms of sample preparation, such as in COAD and READ where roughly a third of germline WXS samples were prepared using WGA. Batch effects present in TCGA data can potentially confound even single cancer type analyses if not properly addressed. In terms of pancancer analysis, the correlation between certain technical factors and cancer types confounds analyses that use cancer type as the phenotype of interest, as we demonstrated in Fig 5. We note that since the initiation of our analysis, the raw TCGA sequence data has moved to the genomic data commons (GDC)[31]. The GDC has realigned the sequence data to the current reference genome (GRCh38 .d1.vd1) using a standardized pipeline to harmonize the BAM file. Although this will eliminate one source of variation (BWA version), it only serves to remind researchers how sensitive data analyses might be to non-standardized data collection protocols, especially in the context of the TCGA data, as our study makes clear.

## Methods

### Cohort

Approval for access to TCGA case sequence and clinical data was obtained from the database of Genotypes and Phenotypes (dbGaP). We selected a total of 9,618 normal tissue DNA samples with whole exome sequence data (S1 Table). We limited analysis to samples sequenced with Illumina technology and aligned to the GRCh37/hg19 reference genome.

### Germline Variant Calling

Aligned sequence data for normal samples in BAM file format and the accompanying metadata was downloaded from CGhub[17]. Individual samples were matched with the target regions for the exome capture kit used to generate the sequence data, and variant calling was limited to these target regions +/− 100 bp. SNVs and small indels were identified using the GATK v.3.5/v.3.4 best practices pipeline and a group calling approach[15,16]. The GATK pipeline includes two preprocessing steps to improve the quality of the BAM file. Local realignment of reads is performed in regions containing indels, and base quality scores are recalibrated to minimize known sources of score bias. 'HaplotypeCaller' was run on individual samples in gVCF output mode, producing an intermediate single sample gVCF to be used for group genotyping. Running this pipeline on a single BAM from CGhub took approximately 15 compute hours and produced a 100MB gVCF. Individual gVCFs were combined in groups of 100 and the final group genotyping step was performed by chromosome on all 9,618 samples as a single cohort. Following this joint genotyping step, all future analysis was limited to the intersection of all exome kit capture regions. The intersection of the kits covered 27 MB and 97.7% of Gencode v19 exons (S2 Table)[18]. GATK VQSR was run separately for SNVs and indels. VQSR learns from variant quality annotations using variants overlapping with vetted resources such as dbSNP and 1000 genomes as a truth set. VQSR filters are defined by the percentage of truth variants that pass filter, termed truth sensitivity (TS). For the initial analysis, SNVs were filtered at VQSR TS 99.5% and indels at VQSR TS 99.0%, as suggested by GATK documentation. PCA was performed on the filtered pancancer VCF using PLINK v1.90b3.29[32]. Multiallelic sites, rare variants (< 1% AF), and sites with missing values were excluded from the analysis, leaving 30,120 variants.

### Annotation and BAM metrics

Putative LOF variants, defined here as stop-gained, nonsense, frameshift, and splice site disrupting, were identified using the LOFTEE plugin for VEP and Ensembl release 85[24]. LOFTEE assigns confidence to loss of function annotations based on position of variant in the transcript, proximity to canonical splice sites, and conservation of the putative LOF allele across primates. For our analysis we used default LOFTEE filter setting and only included high confidence predicted LOF variants. A variant was called LOF if it received a high confidence LOF prediction in any Ensembl transcript.

Predicted variant effects were obtained using Annovar v.2014Jul14[33]. Annovar returns a single prediction for each variant position, collapsing across transcripts and reporting the most damaging variant prediction.

Allele frequencies were obtained from ExAC v0.3.1 and used for comparison to our cohort[23].

We quantified capture efficiency in this analysis as the percentage of capture target area covered by at least 20 X read depth (denoted C20X). Sequence depth information was obtained on BAMs downloaded from CGhub using GATK 'DepthOfCoverage' and the corresponding exon capture bed file to define coverage intervals.

### Realignment Comparison

To assess the effect of heterogeneous alignment protocols on variant calls, we realigned the raw sequence data for a subset of our cohort. We chose 345 samples to represent a large range of sample preparation variation present in the TCGA BAM files (S3 Fig). Reads were stripped from the BAM to generate a FASTQ file using samtools v.0.1.18 bam2fq[34]. The FASTQ was realigned to GRCh37 using BWA MEM v.0.7.12 (with parameters -t 3 -p -M) and duplicates were marked using Picard v.1.131[35,36]. From this point the realigned BAM file was processed through the same GATK pipeline described above to produce individual gVCFs. To directly compare the effect of realignment, we generated a VCF for the 345 realigned samples (NewAlign) and for the same 345 samples processed without the realignment step (OldAlign). We were unable to run GATK indel VQSR on a cohort of this size, thus we filtered both VCFs with GATK SNV VQSR TS 99.5 and GATK indel hardfilters (settings QD > 2, FS < 200, ReadPosRankSum > −20). We calculated discordance between alignment pipelines as the percent discordant variant calls: 1- (intersection of variant calls/ union of variant calls). Variant calls were matched by position and alternate base, disregarding zygosity.

### Repeated Samples

A subset of individuals in our cohort have multiple germline DNA WXS samples. This cohort of 9,618 samples represents 9,099 unique individuals; 1012 of the normal WXS samples were obtained from 493 individuals (2–5 samples per individual). The repeated samples all represent germline DNA from the individual, but differ in terms of sample preparation, sequencing, and processing. Percent discordance between repeated samples was calculated as described above. One sample (TCGA-BH-A0BQ) was removed from future analysis due to a high discordance between two high coverage DNA samples. We suspect a sample label mismatch. For association testing, we selected one the sample with the highest coverage that was not whole genome amplified, leaving 9098 samples.

### Indel Filter Methods

To assess different indel filtering methods, indels were extracted from the raw pan-cancer VCF using GATK ‘SelectVariants‘. Multialleleic sites containing both SNPs and indels were included in the indel VCF. Four filter methods were tested on the pan-cancer indel VCF: GATK VQSR TS 90.0, TS 95.0, TS 99.0, and GATK Hardfilter. GATK VQSR and Hardfilter filters were applied using the modules ‘ApplyRecalibration’ and ‘VariantFiltration’ respectively (Hardfilter settings QD > 2, FS < 200, ReadPosRankSum > −20). Indels were additionally identified using Varscan v.2.3.9 (with parameters --p-value 0.1 --strand-filter 1) on BAMs downloaded directly from CGhub with no preprocessing[25]. Single sample indel VCFs were generated using Varscan for all 9618 samples in our cohort.

### Statistical Methods

To detect contribution of technical factors to LOF variant burden Type II ANOVA was performed using the R "car" package[37]. To determine the percent variance explained by technical factors the sum of squared error for each factor was divided by the total sum of squared error. To detect association between germline gene LOF status and cancer type, we used an ‘one vs. rest’ approach. For each cancer type, a binary (‘dummy’) vector was created indicating whether each individual had the given cancer type (1) or another cancer type (0). For sex specific cancers, only individuals of the same gender were compared. LOF variants with AF < 0.05 were binned by individual by gene to generate on individual LOF variant count for each gene. Genes were only included in our analysis if at least two individuals in the cohort had germline LOF variants in the gene. For each cancer type and each gene we used a logistic regression to test association between germline LOF variant burden and cancer type. Our regression model took the form: glm(cancer type indicator ∼ variant burden + race + age). To discover significant gene-cancer type associations we obtained the p value of the β coefficient for the variant burden term and used a Bonferroni cutoff of 1.61 × 10^−7^ to account for multiple testing (31 cancer types × ∼10,000 genes).

## Acknowledgements

All computing was done using the National Resource for Network Biology (NRNB) P41 GM103504.

All primary data was accessed from The Cancer Genome Atlas Research Network (cancergenome.nih.gov).

AB is supported in part by the National Institute Of General Medical Sciences of the National Institutes of Health under Award Number T32GM008666 and the Translational Genomics Research Institute (TGen). NJS and his lab are also supported in part by National Institutes of Health Grant UL1TR001442 (CTSA) (note that the content of this manuscript is solely the responsibility of the authors and does not necessarily represent the official views of the NIH).

